# How cell penetrating peptides behave differently from pore forming peptides: structure and stability of induced transmembrane pores

**DOI:** 10.1101/2023.07.26.550729

**Authors:** Haleh Alimohamadi, Jaime de Anda, Michelle W. Lee, Nathan W. Schmidt, Taraknath Mandal, Gerard C. L. Wong

**Affiliations:** Department of Bioengineering, University of California, Los Angeles, Los Angeles, CA 90025, USA; Department of Chemistry and Biochemistry, University of California, Los Angeles, CA, 90095, USA; Department of Microbiology, Immunology, and Molecular Genetics, University of California, Los Angeles, CA, 90095, USA; California NanoSystems Institute, University of California, Los Angeles, CA, 90095, USA; Department of Physics, Indian Institute of Technology Kanpur, Kanpur, 208016, India

## Abstract

Peptide induced trans-membrane pore formation is commonplace in biology. Examples of transmembrane pores include pores formed by antimicrobial peptides (AMPs) and cell penetrating peptides (CPPs) in bacterial membranes and eukaryotic membranes, respectively. In general, however, transmembrane pore formation depends on peptide sequences, lipid compositions and intensive thermodynamic variables and is difficult to observe directly under realistic solution conditions, with structures that are challenging to measure directly. In contrast, the structure and phase behavior of peptide-lipid systems are relatively straightforward to map out experimentally for a broad range of conditions. Cubic phases are often observed in systems involving pore forming peptides; however, it is not clear how the structural tendency to induce negative Gaussian curvature (NGC) in such phases is quantitatively related to the geometry of biological pores. Here, we leverage the theory of anisotropic inclusions and devise a facile method to estimate transmembrane pore sizes from geometric parameters of cubic phases measured from small angle X-ray scattering (SAXS) and show that such estimates compare well with known pore sizes. Moreover, our model suggests that whereas AMPs can induce stable transmembrane pores for membranes with a broad range of conditions, pores formed by CPPs are highly labile, consistent with atomistic simulations.

## Introduction

Transmembrane pore formation is central to biological processes critical to the maintenance of complex organisms, and dysregulation of such processes results in a broad range of disease states. For example, peptides such as antimicrobial peptides (AMPs) can bind to bacterial membranes and disrupt the membranes through transmembrane pore formation, blebs, and vesiculation, which can lead to membrane depolarization, leakage, and cell lysis ^1–4^. Bcl-2 family proteins integrate multiple surface stresses, such as hydrophobic insertion and oligomerization into putative supramolecular pores, to govern the intrinsic apoptotic pathway ^5^. What’s more, recent work has shown that histones and fragments thereof can lead to poration processes that precipitate smooth muscle cell death and inflammatory responses in atherosclerosis ^6^. The membrane remodeling necessary for such pore formation is often accomplished via proteins or peptides, which adsorb or insert onto a cell membrane and thereby influence its geometry.

Direct experimental measurements of pore geometry have been a challenge for decades. Several methods have been proposed to measure the radius of transmembrane pores in situ or reconstituted systems, each with its own compromises. These include variations of dye leakage, atomic force microscopy (AFM), electron microscopy (EM), and small-angle X-ray scattering (SAXS) ^2,7,8^. Even with the availability of a structural model, the general expectation is that it is not one single static structure because the formation of transmembrane pores in realistic biological contexts depends on protein or peptide sequence, lipid composition and a broad range of intensive thermodynamic variables. Additionally, the size of pores in the membrane typically varies within the order of nanometers and changes with time and thermal fluctuations, which can render them challenging to measure experimentally. In electron-based structural probes, this difficulty is exacerbated by the potential for radiation damage, which can influence the equilibration of the system, and the intrinsic low signal-to-noise ratio, which can limit contrast in biological samples ^9^.

SAXS is, in principle, a non-destructive technique that has been used extensively to study membrane remodeling at the nanoscale. Using SAXS experiments, it has been previously shown that various AMPs and cell-penetrating peptides (CPPs) can restructure small unilamellar vesicles (SUVs) into bicontinuous cubic phases, which are rich in negative Gaussian curvature (NGC) ^10–18^. In a more general compass, NGC is required to allow communication through a membrane: it has been shown that cubic lipid matrices can be used to promote efficient siRNA delivery ^19,20^. This observed behavior is, in principle, consistent with their biological membrane remodeling activity since NGC or saddle splay curvature is topologically required for the formation of transmembrane pores. However, it is not clear how to correlate the measured induced curvatures in the cubic phases (such as the NGC density per volume) to the actual size of transmembrane pores in the membrane (Fig. 1). Indeed, cubic phases are idealized as minimal surfaces with zero mean curvature, so that the principal curvatures c_1_ and c_2_ are equal in magnitude but opposite in sign at every point (Fig. 1), a condition that is not met (a least in general) for typical transmembrane pores which have extremely high positive curvature c_1_ (∼1/2 nm^-1^).

**Figure 1.**
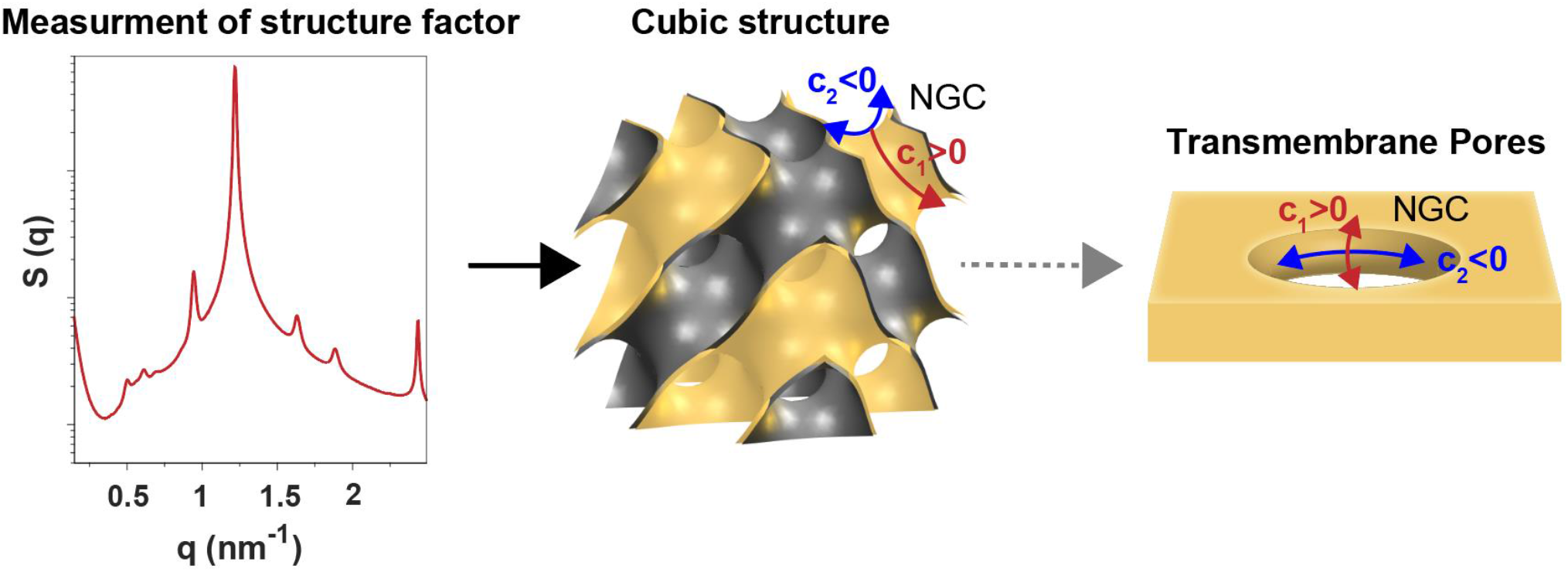
Using induced membrane curvature deformations by NGC-generating inclusions in SAXS spectra to estimate the radius of transmembrane pores generated by pore forming peptides. Cubic structure and transmembrane pores are both characterized by a saddle shape (NGC) with positive and negative principal curvatures (c_1_ > 0 and c_2_ < 0).

In recent years, discrete and continuum theoretical approaches combined with information from experiments have emerged as a powerful method to study the induced membrane curvatures by proteins and peptides ^21–29^. Discrete models are powerful for exploring molecular mechanisms underlying lipid-peptide interactions. In contrast, continuum models work best when engaging membrane phenomena at large scales, such as processes that require membrane deformations over hundreds of nanometers ^30–34^. For example, Akimov et al. applied continuum elasticity theory to analyze the energy landscape of pore formation and closure ^35,36^. They proposed that the magnitude of line tension at the edge of a transmembrane pore depends on the radius of the pore and external surface stresses ^35,36^. Betterton et al. studied the competition between electrostatics effects and line tension in opening a transmembrane pore and later Fošnarič et al. showed the ability of anisotropic inclusions to stabilize transmembrane pores in the membrane ^37,38^. However, it is not clear how to generalize these methods to estimate pore sizes from SAXS experiments for different protein/peptide sequences, pH conditions, or temperatures.

In this work, we present a minimal mechanical model at the continuum length scale to estimate the radius of peptide-induced transmembrane pores based on peptide-induced membrane deformations measured in SAXS experiments conducted specifically for different peptide sequences, temperature, or other intensive thermodynamic variables (Fig. 1). Specifically, we estimate the magnitude of induced curvatures by peptides from measured lattice constants of cubic phases and minimize the excess free energy of transmembrane pore formation by leveraging a model based on anisotropic membrane inclusions.

We systematically explore the influence of the cubic lattice constant, peptide surface area, electrostatic repulsion, and interfacial tension at the pore’s edge on the opening and closure of transmembrane pores. The model indicates that the pore radius is a non-monotonic function of the cubic lattice constant; the pore radius initially increases with the cubic phase lattice constant, but eventually saturates and then decreases. We show that the pore radii calculated with our model are in agreement with the experimental measurements. Furthermore, this mechanical framework is used to estimate the sizes of pores generated by AMPs and CPPs and analyze the stability of these induced pores to variations in membrane properties. Interestingly, we find that transmembrane pores induced by CPPs are intrinsically much more labile than those by AMPs when membrane perturbations are considered. These results are consistent with those from atomistic simulations, and independent of effects from peptide residence time in the membrane. We believe our proposed framework is generalizable to other pore forming systems and can potentially provide insight into the development of new antimicrobials and drug delivery systems.

## Model development

### Membrane mechanics

Our system consists of a lipid membrane and NGC-generating inclusions (proteins or peptides) confined in the saddle-shaped domain at the rim of transmembrane pores. Previous studies have shown that AMPs have a stronger binding affinity for the pore domain in order to stabilize pore formation ^39^. In this study, we model the lipid bilayer as a continuous elastic shell with negligible thickness and assume that the membrane is areally incompressible ^40,41^. This allows us to simplify the total free energy of the system (*E*_*total*_) as ^37,38,42–44^

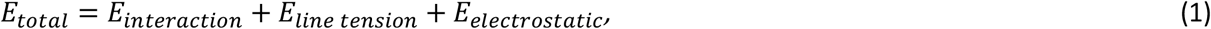

where *E*_*interaction*_ is the elastic energy due to the mismatch between the local shape of the lipid membrane and the intrinsic shape of the inclusion, *E*_*line tension*_ is the energy cost due to the exposure of the hydrophobic tails of lipid molecules at the edge of the transmembrane pore, and *E*_*electrostatic*_ is the electrostatic repulsion energy between the charged lipids around the pore. Assuming that the membrane inclusions are more rigid than the lipid bilayer and impose their intrinsic curvatures on the underlying membrane ^45^, the membrane-inclusion interactions energy is given as ^42–44^

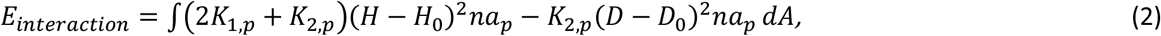

where *K*_1,*p*_ and *K*_2,*p*_ are constants (*K*_1,*p*_> 0 and *K*_2,*p*_< 0) representing the strength of interaction between the inclusions and the surrounding membrane ^42,43^. *H* is the mean curvature of the membrane and *D* denotes the curvature deviator of the lipid membrane (*H*^2^ − *D*^2^ =*K* is the Gaussian curvature). *H*_0_ and *D*_0_ are the intrinsic mean and deviatoric curvatures of the inclusion, respectively. *a*_*P*_ is the area of the inclusion in contact with the surrounding membrane, and *n*is the density of the inclusions on the membrane surface. In Eq. 2, we assume that attached inclusions have a homogenous lateral distribution on the surface and impose their intrinsic curvatures on the underlying membrane ^42^ and the integral is taken over the area of the membrane that is covered by NGC-generating inclusions.

The interfacial energy at the edge of the transmembrane pore in Eq. 1 is given as

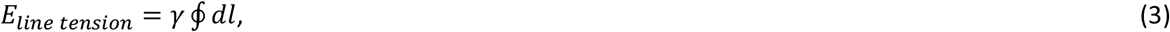

where *γ* is a constant representing the magnitude of the interfacial force and the integral is taken over the interfacial line *∂l*. The line tension energy favors closing the transmembrane pores in lipid bilayers. The magnitude of line tension (*γ*) depends on various factors, such as the lipid composition ^46–49^, the radius of the pore ^35^, the interaction between the membrane and inclusions such as AMPs ^50^, and external stresses applied to the membrane ^36^. Here, for simplicity, we assume that the line tension is constant and independent of the lipid composition or membrane morphology.

We model the electrostatic repulsion energy in Eq. 1 using the linearized Poisson-Boltzmann formalism for thin membranes, which is given by ^37,38,51,52^

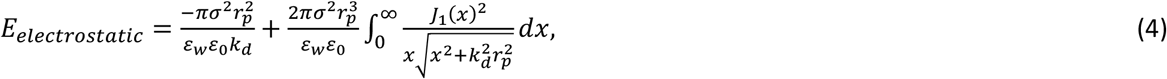

where *σ* is the surface charge density, *r*_*p*_ is the radius of a pore, *ε*_0_ is the permittivity of free space, *ε*_*w*_ ≈ 80 is the dielectric constant of the aqueous solution, *K*_*d*_ is the inverse of Debye length 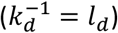,and *J*_1_ is the Bessel function. The electrostatic energy favors the opening and enlargement of transmembrane pores.

To establish the relationship between the induced curvatures by pore former peptides observed in SAXS experiments to the radius of the stable transmembrane pores, we assume that pore former peptides generate a single circular pore with a radius of *r*_*p*_ on a planar membrane. We model the transmembrane pore as a semitoroidal cap with a height equal to the bilayer thickness (2*b*, Fig. 2A), and write the excess free energy with respect to a flat membrane with no pore as

**Figure 2.**
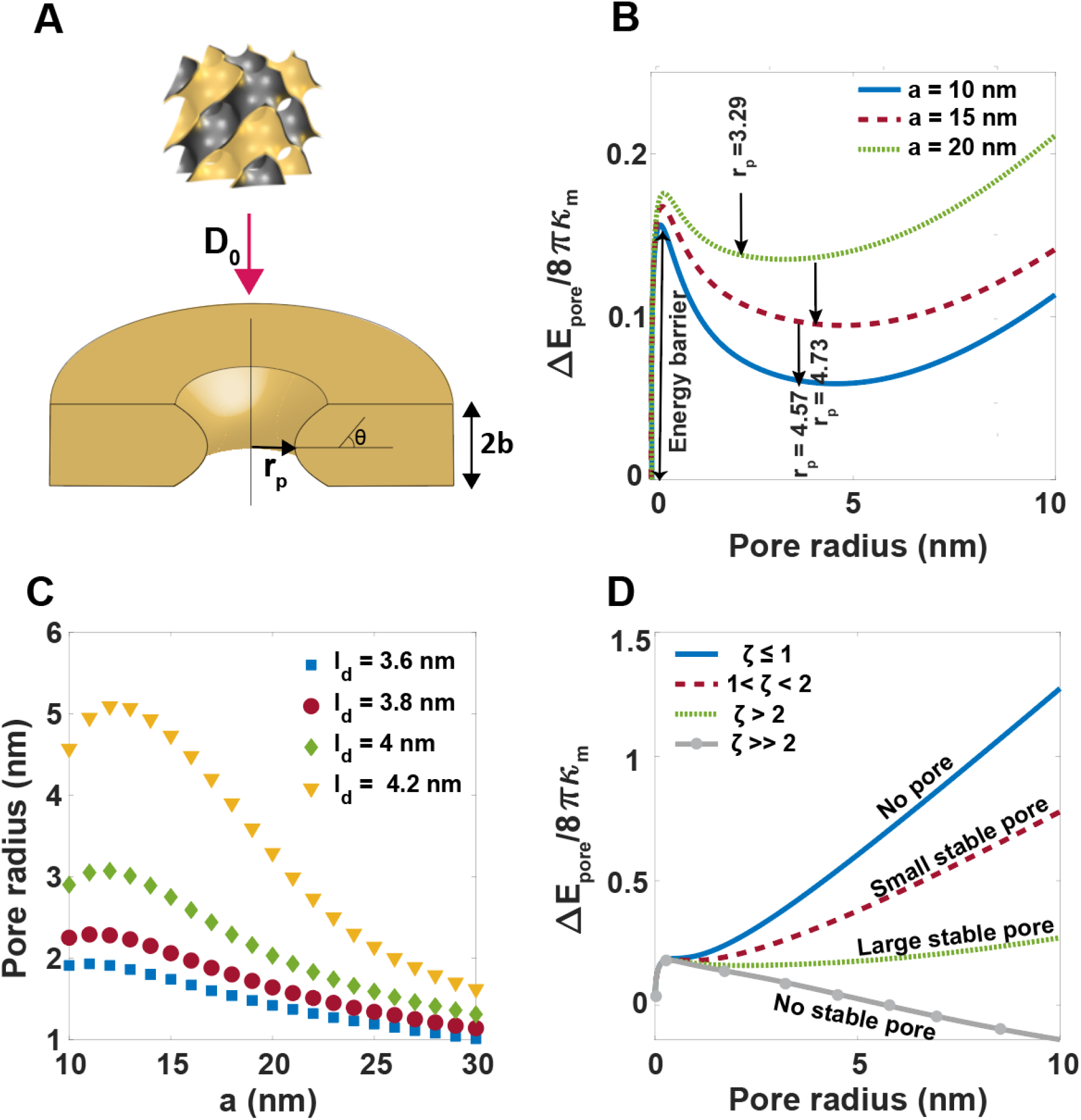
Estimating the radius of transmembrane pores induced by pore former peptides based on the generated cubic structures in SAXS experiments. **(A)** The induced deviatoric curvature (*D*_0_) in a cubic structure by a pore former peptide is used to estimate the radius of a circular pore in a planar membrane. The transmembrane pore is modeled as a semitoroidal cap with a radius of r_p_ and a height equal to the bilayer thickness 2*b*(π/2 < θ *<* 3π/2). (**B**) The change in the energy of the system from a planar membrane with no pore to a planar membrane with a single circular pore (Eq. 5) as the function of the pore radius for three different lattice constants (a*p* = 4 nm^2^σ = -0.05 As/m^2^, and *ld* = 4.2 nm, γ = 10 pN, and using *Pn3m*). Arrows show the location of minimum energy that corresponds to the radius of a stable transmembrane pore formed in the membrane. (**C**) The non-monotonic function of pore radius with increasing the cubic lattice constant (a_*p*_ = 4 nm^2^, σ = -0.05 As/m^2^, and γ = 10 pN). (**D**) The change in the energy of the system as a function of pore radius for different ratios of Debye length to the characteristic length of peptide, 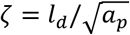, σ = -0.05 As/m^2^, and γ = 10 pN).

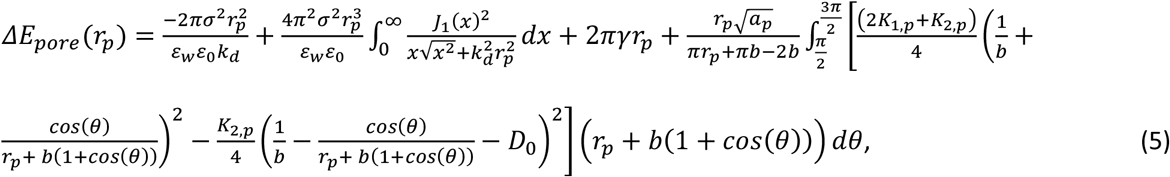

where *θ* is the angle between the vertical axis and the boundary of the semitoroidal cap. It should be mentioned in Eq. 5, we assume that the induced mean curvature by NGC-generating inclusions is negligible compared to the induced deviatoric curvature (*H*_0_≪ *D*_0_) and also restrict the number of inclusions in the rim region based on the steric repulsions between inclusions 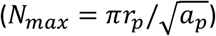.

We estimate the magnitude of the induced deviatoric curvature (*D*_0_ in Eq. 5) by NGC-generating inclusions based on the identified cubic structures in SAXS measurements given as ^53,54^

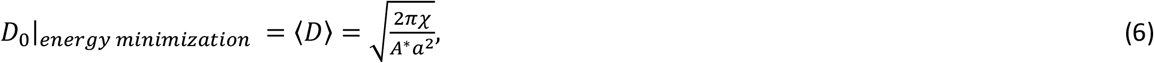

where *a* is the lattice parameter, *χ* is the Euler characteristic, and *A*^∗^ is the surface area per unit cell specific to each cubic phase ^53,54^. Using SAXS data, we can determine the type of the cubic phases and their corresponding lattice constants, which allows us to calculate the magnitude of induced curvatures (Figs. 2A and 3A). Assuming the total area of the membrane remains constant, we numerically calculate the excess energy of the system (Eq 5) and find the pore radius (*r*_*p*_) that minimizes this energy. We also estimate the strength of interaction between the membrane and inclusions using the elastic properties of the membrane and the characteristic length of the inclusions (Eqs. S10-S12).

## Results and Discussion

### Estimation of the radius of transmembrane pores generated by pore former peptides

A range of peptides can remodel lipid membranes and open transmembrane pores. The size of transmembrane pores induced by peptides is a key geometric parameter in mediating different processes such as apoptosis, cell death, and pathogen internalization ^16,55^. Here, we use our mechanical framework to estimate the radius of transmembrane pores induced by pore former peptides from SAXS measurements, and systematically explore the influence of cubic lattice constant, projected peptide surface area on the membrane, interfacial tension, and charge density on the size of the generated pores.

We used the deviatoric curvature (*D*_0_) directly extracted from lattice constants of *Pn3m* cubic structures measured in SAXS as input to a planar membrane with a semi-toroidal pore (Fig. 2A). We minimized the energy of the system as described in Eq.5. The change in the energy has a local minimum which represents the formation of a stable semi-toroidal pore in the membrane with the presence of NGC-generating inclusions (Fig. 2B, *a*_*p*_= 4 nm^2^, σ = -0.05 As/m^2^, *l*_*d*_ = 5 nm, and γ = 10 pN). We observed that pore formation in the membrane is associated with an energy barrier and the height of the energy barrier increases as the lattice constant of the cubic phase decreases (Fig. 2B). This energy barrier characterizes the transition from a hydrophobic defect into a hydrophilic pore ^35,56^. For membranes with no inclusions, previous studies have shown that the local minimum of the energy for transmembrane pore formation is shallow (below *KT*) and the pore likely closes from thermal fluctuations ^37,38^. However, partitioning of anisotropic inclusions such as pore forming peptides (e.g., AMPs) in the membrane rim can significantly lower and deepen the local minimum of the energy, which leads to the formation of stable and long-lasting transmembrane pores ^16^.

It is interesting to note that the radius of the pore is a non-monotonic function of the cubic lattice constant. As the cubic lattice constant increases, the radius of the pore increases, saturates, and then decreases (Fig. 2C, *a*_*p*_= 4 nm^2^, σ = -0.05 As/m^2^, and γ = 10 pN). This can be due to the different distribution of anisotropic curvature in a semi-toroidal pore versus a cubic phase: The principal curvatures within a transmembrane pore are strong anisotropic: A large positive curvature is necessary for the lip that connects the outer leaflet with the inner leaflet, whereas a range of negative curvatures can be accommodated in the circumference of the pore. This contrasts with curvatures in a cubic phase, where the positive and negative curvatures are constrained to be equal in magnitude. Moreover, our results showed that the non-monotonic relationship between the pore radius and the cubic lattice constant becomes even more prominent with an increase in Debye length (Fig. 2C) and surface charge density (Fig. S1A), and a decrease in the projected surface area of the peptide in contact with membrane (Fig. S1B) and line tension (Fig. S1C).

We identified three different regimes of pore forming behavior, depending on the ratio of the Debye length to the characteristic length of the pore former peptide, 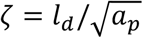. In Fig. 2D, we plotted the change in the energy of the membrane as a function of the pore radius for a fixed lattice constant (*a* = 25 nm, σ = -0.05 As/m^2^, and γ = 10 pN). (I) At small ζ (ζ ≤ 1 nm), the change in energy increases with increasing the radius of the pore, indicating that the holes always tend to close (solid blue line in Fig. 2D). (II) At intermediate ζ (1 < ζ < 2 nm), the energy is minimized at a small pore radius, *r*_*p*_∼ 1 nm (dashed red line in Fig. 2D). (III) At large ζ (ζ > 2 nm), the change in energy is minimized for a large pore radius, *r*_*p*_> 2 nm (dotted green line in Fig. 2D). This suggests that pore former peptides can generate large and stable holes in lipid membranes through conformational changes, such as adopting β-strand structures^57^, to reduce the peptide-membrane contact surface area. For peptides with ζ ≫ 2 nm, the change in the energy decreases linearly with increasing pore radius (gray line with circles in Fig. 2D). Therefore, hole radii grow in the membrane and there is no stable transmembrane pore (Fig. 2D).

### The transmembrane pore formation exists over a range of cubic lattice constants, Debye length, surface charge density, line tension, and peptide surface area

Over what ranges of cubic lattice constants, surface charge density, and line tension, peptides with different sizes can open a stable transmembrane pore? To answer this question, we performed numerical calculations and calculated the radius of a stable transmembrane pore for a range of measured lattice constants in SAXS experiments, reported surface charge density and Debye length for lipid membranes, as well as over a range of hydrophobic line tensions and projected surface area of peptides. The results are summarized in a series of ‘phase diagrams’ (Figs. 3A-3D). The gray and white regions in Figs. 3A-3D represent the areas with no pore and no stable pore formation, respectively.

**Figure 3.**
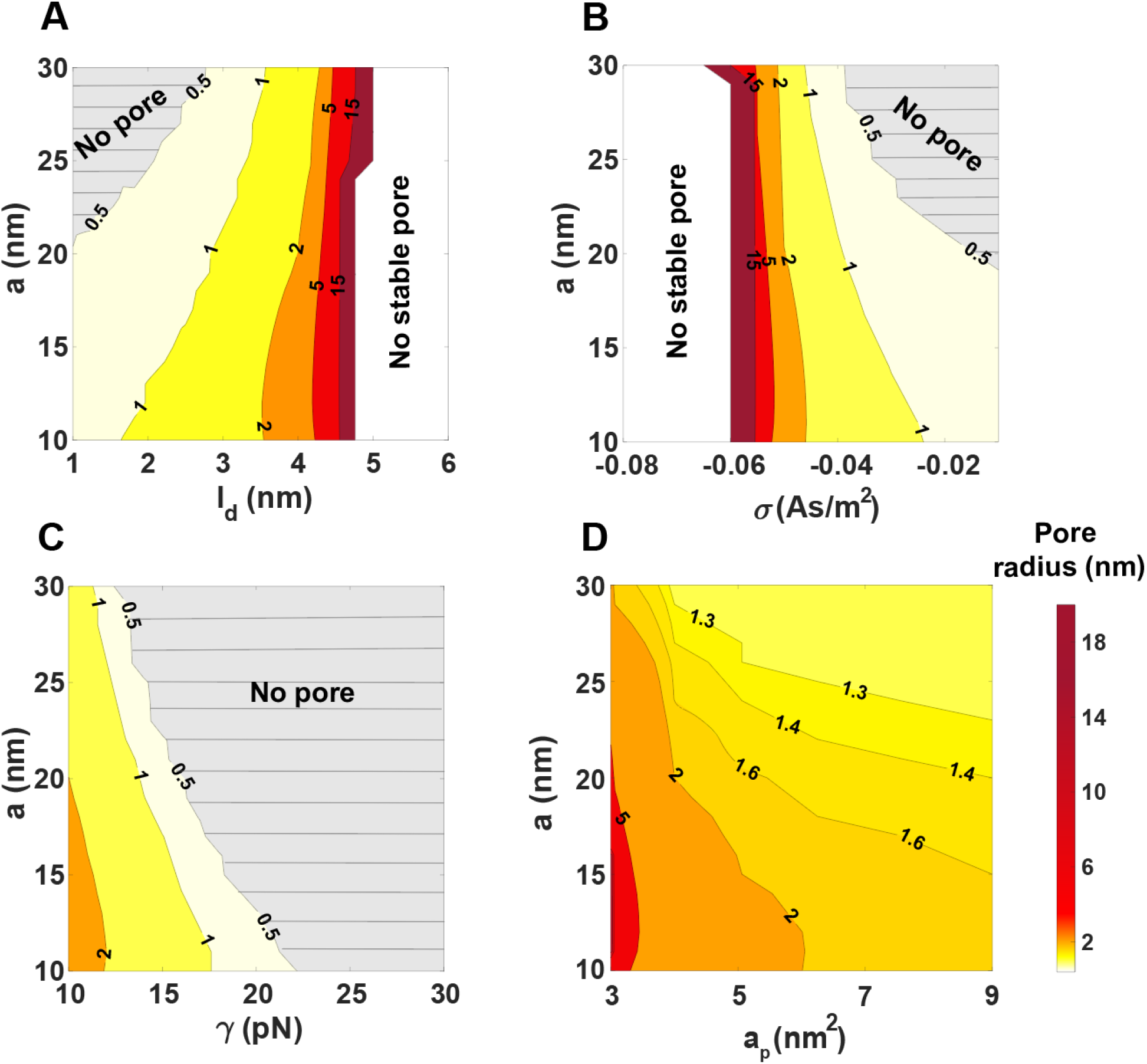
The size of transmembrane pores depends on the cubic lattice constant (*a*), charge density (*σ*), line tension (γ), and the surface area of the peptide (*a*_*p*_). Contour plot of the transmembrane pore radius for a range of (**A**) cubic lattice constants and Debye length (σ = -0.05 As/m^2^, a*p* = 4 nm^2^, and γ = 10 pN), (**B**) cubic lattice constants and charge density (σ = -0.05 As/m^2^, *ld* = 4 nm, a*p* = 2 nm and γ = 10 pN), (**C**) cubic lattice constants and line tension (σ = -0.05 As/m^2^, a*p* = 4 nm^2^, and *ld* = 4 nm), (**D**) cubic lattice constants and peptide surface area (σ = -0.05 As/m^2^, and *ld* = 4 nm, and γ = 10 pN). The gray domains in panels (**A**), (**B**), (**C**), and (**D**) mark the regions with no pores and the white domains represent the region with no stable pores.

For typical properties of lipid bilayers, the Debye length and surface charge density vary in a wide range between 1 < *l*_*d*_ < 10 nm ^58,59^ and −0.0075< σ <-0.08 As/m^2^^51^, respectively. As expected from Eq. (4), with increasing the Debye length and the magnitude of surface charge density, the electrostatic repulsion becomes dominant, which results in the formation of larger pores in the membrane (Figs. 3A and 3B). Another factor that controls the size of the pore is induced line tension at the edge of the pore. The magnitude of line tension varies widely between 10 < γ < 200 pN, depending on the lipid composition, particularly the molar fraction of cholesterol ^47,60,61^. Our results show that line tension significantly reduces the size of transmembrane pores. For example, for fixed σ = -0.05 As/m^2^, *a*_*p*_= 4 nm^2^, and *l*_*d*_ = 4 nm, no pore can be formed in the membrane when *γ* > 20 pN, regardless of the magnitude of the cubic lattice constant (Fig. 3C).

Previous studies have shown that the structure and conformational stability of pore forming peptides are key factors determining their antimicrobial activity ^62–64^. To explore the relationship between the peptide structure and its antimicrobial activity, we estimated the induced pore sizes for a range of peptide contact surface areas with the membrane and cubic lattice constants (Fig. 3D). The projected surface area of the peptides affects the strength of the membrane-peptides interactions (Eq. S10), and it also characterizes the number of peptides that can fit within the membrane rim due to steric repulsions ^16^. We observed that the pore radius decreases continuously with an increase in the projected surface area of the peptide (Fig. 3D). Overall, our results provide important insights into the synergistic role of electrostatic repulsions, interfacial forces, and induced curvatures by peptides in controlling the formation of stable pores in the membrane.

### Membrane remodeling behavior of AMPs and CPPs: numerical comparisons of robustness of induced transmembrane pores using continuum framework

Having established a mathematical model to estimate the size of membrane holes generated by pore former peptides from the induced cubic structures in SAXS experiments, we applied the framework to explore (*I*) the range of pore sizes induced by AMPs and CPPs and (*II*) robustness of the formed pores against variations in the membrane properties. In our previous work, we utilized binary lipid mixtures (PS/PE 20/80 or PG/PE 20/80) and measured the lattice constant of cubic phases induced by AMPs including melittin, magainin (Fig. S5), protegrin-1 (PG-1), human-β-defensin-2 (HBD-2), rhesus-myeloid-α-defensin-4 (RMAD-4), HBD-2, mouse-α-defensin 4 (CRP-4), and CPPs such as HIV TAT and R-9 ^13,10,14–16^. Using the reported cubic lattice constants and estimating the effective surface area of pore former peptides based on their amino acid sequence and structures, we calculated the radius of membrane pores generated by a variety of AMPs and CPPs for fixed *κ*_*d*_ = 4 nm^-1^, σ = -0.05 As/m^2^, and γ = 10 pN (Table S2).

Based on our results, AMPs can induce stable transmembrane pores with a radius between 1-3 nm, which is in agreement with experimental measurements of pore sizes (Table S2) ^22,28,65–69^. However, our model predicts that CPPs, R-9 and HIV TAT, induce pores that are not stable in the membrane (Table. S2). Previous experimental studies have also suggested that CPPs such as R-9 can only generate transient pores, and HIV TAT peptides are only capable of inducing pores with high concentration and in model membranes that contain a significant proportion of anionic lipids or lipids that induce negative intrinsic curvatures ^65,69^. Similarly, it has been suggested that cell penetrating peptide pep-1 translocates across the cell membranes without forming a stable pore ^22,28,66,67^. It is not clear, however; whether these labile pores are a consequence of expected shorter membrane residence times of cell penetrating peptides, which typically have much less hydrophobicity than AMPs.

To investigate the intrinsic stability of the induced transmembrane pores of a specific pore size against variation in membrane properties, we introduced a 5% perturbation in the inverse of Debye length (*κ*_*d*_), the surface charge density (σ), the line tension (*γ*), and then calculated the resulting pore radius (Fig. 4). Interestingly, we found that AMPs display a high degree of stability in generating membrane pores across a range of membrane properties (Fig. 4). With 5% variation in membrane properties, we observed that the radius of induced pores by AMPs, except for PG-1 and magainin, changes within a narrow range between 1-3 nm. PG-1 and magainin generate pores with a wider range of radii (between 1-5 nm), particularly in response to perturbations in *κ*_*d*_ and σ (Figs. 4A and 4B). The formation of large, stable pores (with a diameter of 9 nm) by PG-1 was also demonstrated in a previous study using atomic force microscopy ^22^.

**Figure 4.**
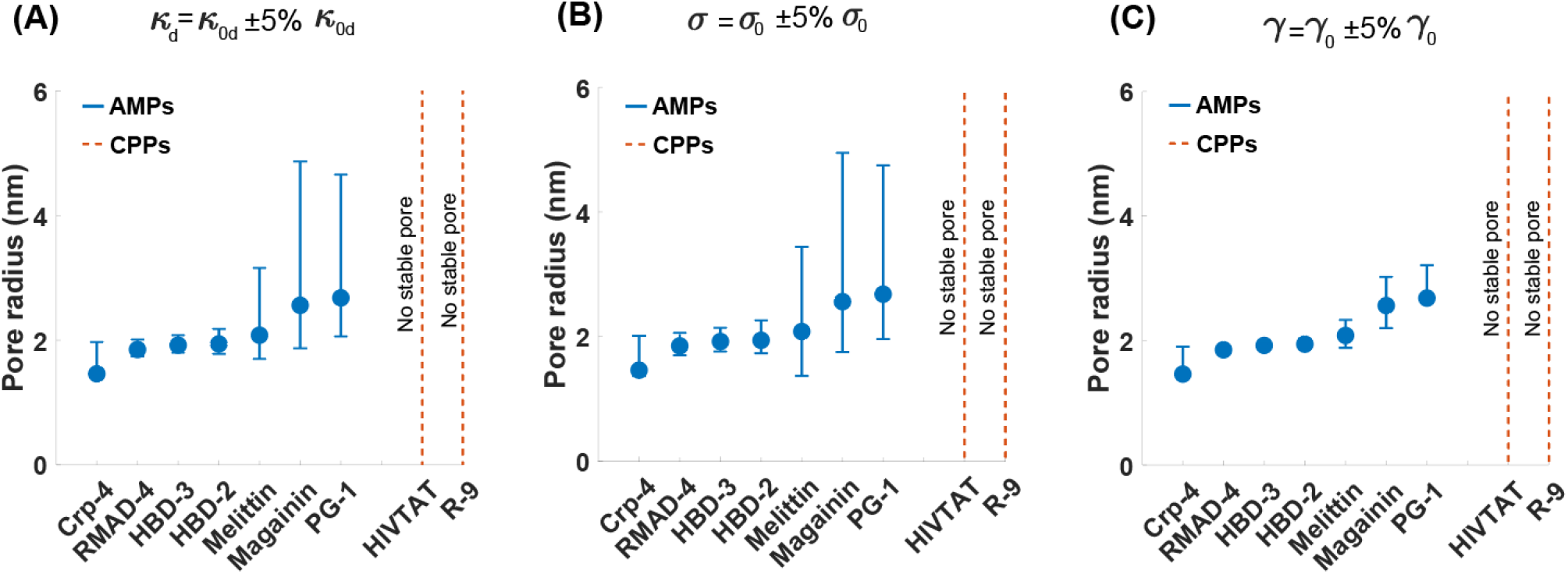
Sensitivity analysis of pore sizes induced by AMPs and CPPs to variation in the membrane properties. We used the measured cubic lattice constants of AMPs ^13–16^ and CPPs^10^ in our previous studies and calculated the pore radius (Eq. 5) with 5% variation in (**A**) the inverse of Debye length (*κ*_*d*_), (**B**) the surface charge density (σ), and (**C**) the line tension (γ). *κ*_0d_ = 4 nm^-1^, σ_0_= -0.05 As/m^2^, and γ_*0*_ = 10 pN. AMPs are robust in generating stable pores with variations in membrane properties.

In contrast to AMPs, our results indicate that formed ‘pores’ by CPPs are highly unstable with respect to perturbations in environmental parameters such as membrane surface charge density, Debye length, line tension (Fig. 4). We hypothesize that such instability is intrinsic to small radii pores induced by CPPs, due to the strong elastic restoring forces from high local membrane curvature deformations. This instability makes measuring the accurate size of formed pores by CPPs challenging, considering that CPPs usually induce small transient pores in the membrane ^65,69^. Thus, our framework predicts a transition in the character of the pores as the pore radius decreases, from large stable pores from AMPs to labile pores from CPPs. More generally, from a methodological perspective, we have presented a minimal theoretical framework for *(I)* estimating pore size from the structure of induced cubic phases in SAXS, *(II)* analyzing the stability of pores against perturbations in the lipid bilayer properties, and *(III)* designing a reconstituted system in order to induce a desired pore size with a given pore former peptide or protein.

### A comparison of the stability of transmembrane pores induced by AMPs and by CPPs, using atomistic molecular dynamics simulations

Our results from the continuum framework suggest that AMPs but not CPPs have the ability to induce stable transmembrane pores. To test this hypothesis, we conducted fully atomistic Molecular Dynamics (MD) simulations using both AMPs and CPPs. Previous studies have revealed spontaneous pore formation in membranes by AMPs is challenging to achieve in fully atomistic simulations because, as we have shown in Fig. 2, pore formation is associated with a high energy barrier ^27^. Therefore, we investigate if AMPs and CPPs can stabilize a preformed membrane pore, which, in the absence of any peptide, undergoes spontaneous closure. We first constructed a transmembrane pore with five magainin AMPs in a DOPS/ DOPE (20/80) bilayer (Fig. 5A). Then, the bilayer was equilibrated through molecular dynamics (MD) simulations (see atomistic MD method section in the supplement).

**Figure 5.**
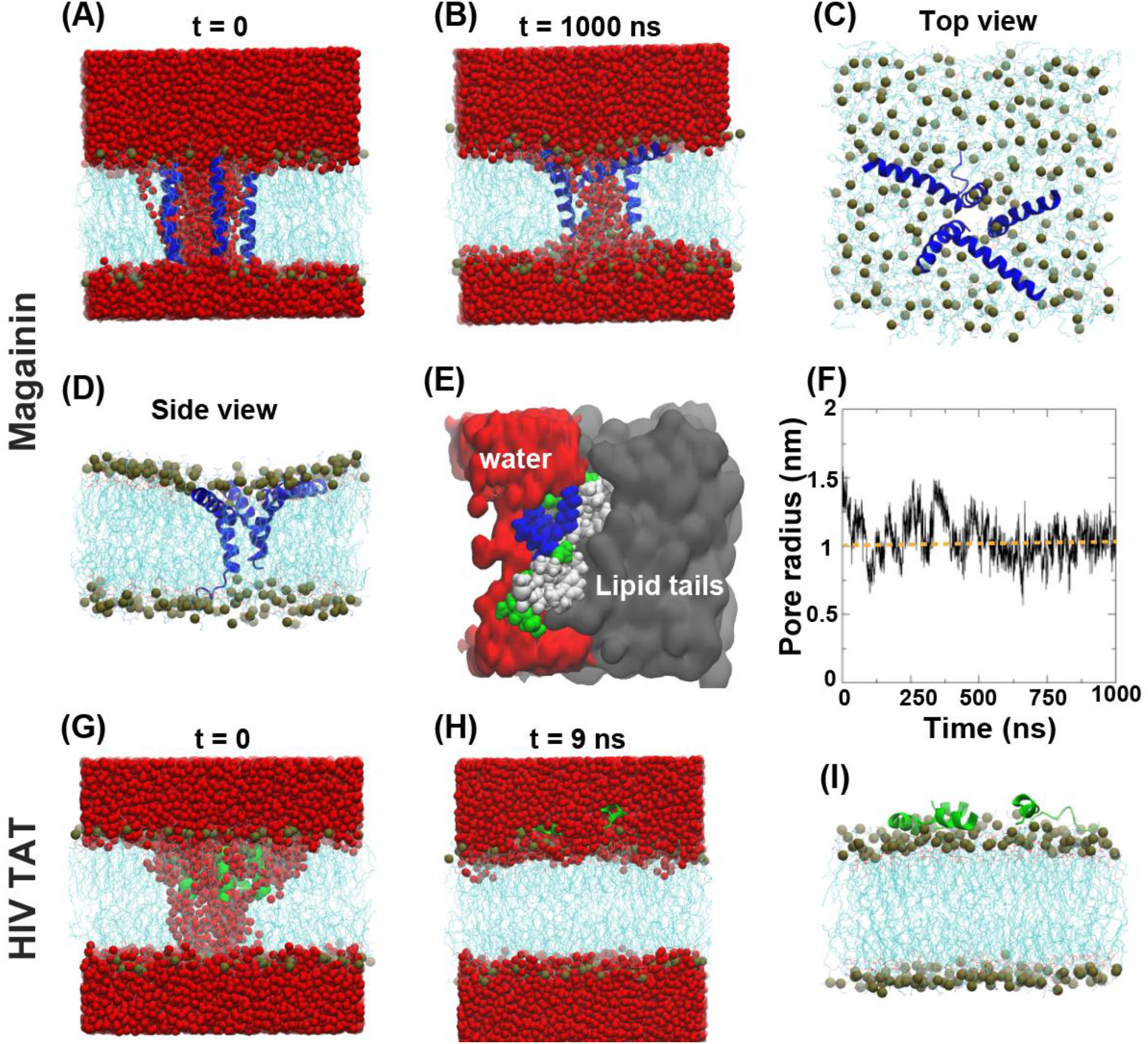
Molecular dynamics simulations to investigate the pore stabilization capability of magainin AMP versus HIV-TAT CPP. **(A)** Structure of the preformed transmembrane pore with five magainin peptides at the beginning of the simulation. (**B**) Equilibrated structure of the transmembrane pore after 1000 ns long MD simulation. (**C**) Top and (**D**) side view of the equilibrated membrane pore without water molecules. Blue and cyan colors represent peptides and lipids, respectively. Oxygen atoms of the water molecules are shown by red spheres. Bile spheres represent the phosphorous atoms of the lipid heads. (**E**) A zoomed-in image of the pore structure shows that the hydrophobic residues (white) bind with the lipid tails (gray) and the hydrophilic residues (blue and green) are in contact with the water (red). Lipids and water are shown in the ‘surf’ representation. (**F**) The radius of the pore as a function of simulation time. (**G**) Structure of the preformed transmembrane pore with five HIV TAT peptides at the beginning of the simulation. (**H**) The membrane pore closes within only 9 ns of equilibration. (**I**) Side view of the membrane after the closure of the pore, the peptides move out from the membrane core. Green and cyan colors represent the HIV TAT peptide and the lipids, respectively.

Our results show that embedded magainin peptides spontaneously readjusted their positions during equilibration to form a stable transmembrane pore (Fig 5B). Three out of the five peptides lined up along the water channel, and the remaining two adopted a tilted configuration near the pore edge, forming a semi-toroidal shaped transmembrane pore (Figs. 5C and 5D). The amphipathic characteristics of the AMPs contribute to stabilizing the membrane pore, where the hydrophobic residues are in contact with the hydrophobic core of the membrane, and the hydrophilic residues face the water channel (Fig 5E). We also calculated the radius of the pore as a function of simulation time to gain insights into the dynamics of the pore (Fig 5F). Based on our results, the water channel remained stable throughout the 1000 ns long MD simulation, and the radius of the pore was ∼1 nm, which is in good agreement with experimental data ^2,66,67^. We also performed a control simulation by removing the magainin peptides from the preformed membrane pore and equilibrating the structure. We observed that the pore closed immediately within only 2 ns, confirming the role of magainin AMPs in stabilizing the transmembrane pore (Fig. S3).

Next, we repeated the simulations with five HIV TAT CPP, following the same protocol described earlier (Fig. 5G). In contrast to AMPs, we observed that the transmembrane pore was not stable (Fig. 5H). The peptides moved out from the membrane core, resulting in the closure of the pore within a short time of 9 ns (Figs. 5H and 5I). This finding suggests that HIV TAT peptides can generate transient pore morphologies probably due to their strong interactions with the anionic lipids ^69^. However, they are unable to stabilize the induced pore due to lack of hydrophobic residues that can bind with the lipid tails. Our MD simulation findings is in agreement with the experimental study by Kubitscheck et al. which reported the accumulation of HIV TAT peptides on an anionic GUV, leading to transient pore formation and rapid translocation of the peptides across the membrane. However, the lifetime of the generated transmembrane pores was too short to be detected by single-molecule tracking ^69^. Taken together, our molecular dynamic simulations and theoretical results suggest that AMPs are more robust than CPPs in generating and stabilizing the process of transmembrane pore formation.

In summary, by constructing a minimal mechanical model combined with information from SAXS measurements, we are able to estimate the radius of transmembrane pores. We systematically investigate the synergetic effects of the peptide surface area in contact with the membrane, the magnitude of curvatures induced by peptides, and the physical properties of the lipid membrane in governing the size of transmembrane pores (Figs. 2 and 3). By leveraging SAXS measurements across different lipid compositions and pH, we are able to examine the fundamental determinants of pore formation from a new perspective. We find that the size of a transmembrane pore is a non-monotonic function of the cubic lattice constant; as the lattice constant of the cubic phase increases, the pore radius increases, saturates, and then decreases (Fig. 2).

We also identify the range of lipid membrane physical properties combined with the projected surface area of peptides and induced curvatures to generate stable transmembrane pores (Fig. 3). Additionally, we find that pores induced by AMPs are drastically more stable than those induced by CPPs, just from the standpoint of curvature elasticity, independent of the membrane residence times of peptides of different hydrophobicity (Fig. 4). Atomic molecular dynamic simulations validated our predictions that AMPs but not CPPs can stabilize transmembrane pores for a long period of time (Fig. 5). It is interesting to note that CPPs are typically short, with molecular lengths comparable to the elastic decay length of the membrane. Moreover, CPPs typically have fewer hydrophobic residues that insert into hydrophobic cores of membranes compared to AMPs. These observations imply that CPPs will tend to have weak interactions with membrane (Eq. S12). We expect these effects to synergize with the results presented above on the size dependence of pore stability, and thereby render high curvature pores even less stable, as indicated in atomistic simulations. Finally, we anticipate that results here are generalizable to pore former proteins such as viral fusion proteins ^57,70^ and a much broader repertoire of molecules, including synthetic molecules not based on amino acids.

## Methods

The complete derivations with details are given in the Supporting Information.

## Supporting Information

Complete derivations with details; Supplemental Table S2 including the projected surface area of peptides, cubic lattice constant, estimated pore sizes, and the reported pore size induced by each peptide in the literature; Supplemental Figure S1 shows the non-monotonic behavior of pore radius as a function of cubic lattice constant and Figure S2 is the SAXS intensity profile for magainin.

## Acknowledgments

We thank Dennis Bong for the generous gift of magainin peptide. This work was supported by American Heart Association (AHA 966662) grant to G.C.L.W. H.A was supported by the Vascular Biology Training Grant (T32 HL069766-21) and J.D.A. is supported by NSF Graduate Research Fellowship Program (DGE-1650604). T.M gratefully acknowledges the support from the Government of India: Science and Engineering Research Board via Sanction No. SRG/2022/000548. We thank the Stanford Synchrotron Radiation Lightsource (SSRL) (Menlo Park, CA, USA) for access to beamline 4-2. Use of the SSRL, SLAC National Accelerator Laboratory, is supported by the U.S. Department of Energy, Office of Science, Office of Basic Energy Sciences under contract no. DE-AC02-76SF00515. The SSRL Structural Molecular Biology Program is supported by the U.S. Department of Energy, Office of Biological and Environmental Research, and by the National Institutes of Health, National Institute of General Medical Sciences (including P30GM133894). T.M is grateful for the computational resources provided by PARAM Sanganak under the National Supercomputing Mission, Government of India, at the Indian Institute of Technology, Kanpur.

## Author Contributions

H.A designed the research and G.C.L.W supervised the research. T.M conducted the molecular dynamic simulations. H. A, N. W. S, J.D.A, and M. W. L conducted synchrotron SAXS experiments. H.A, J.D.A, M. W. L, T.M, and G.C.L.W wrote and edited the manuscript. All authors reviewed and approved the final version of the manuscript.

